# MyD88-TLR4-dependent choroid plexus activation precedes perilesional inflammation and edema in intracerebral hemorrhage

**DOI:** 10.1101/2022.09.06.506660

**Authors:** Kevin Akeret, Raphael M. Buzzi, Bart R. Thomson, Nina Schwendinger, Jan Klohs, Nadja Schulthess, Livio Baselgia, Kerstin Hansen, Luca Regli, Florence Vallelian, Michael Hugelshofer, Dominik J. Schaer

**Affiliations:** Department of Neurosurgery, Clinical Neuroscience Center, Universitätsspital and University of Zurich, Zurich, Switzerland; Division of Internal Medicine, Universitätsspital and University of Zurich, Zurich, Switzerland; Institute for Biomedical Engineering, University of Zurich and ETH Zurich, Zurich, Switzerland

## Abstract

The functional neurological outcome of patients with intracerebral hemorrhage (ICH) strongly relates to the degree of secondary brain injury (ICH-SBI) evolving within days after the initial bleeding. Different mechanisms including the incitement of inflammatory pathways, dysfunction of the blood–brain barrier (BBB), activation of resident microglia, and an influx of blood-borne immune cells, have been hypothesized to contribute to ICH-SBI. Yet, the spatiotemporal interplay of specific inflammatory processes within different brain compartments has not been sufficiently characterized, limiting potential therapeutic interventions to prevent and treat ICH-SBI. Using a whole-blood injection model in mice, we systematically characterized the spatial and temporal dynamics of inflammatory processes after ICH using 7-Tesla magnetic resonance imaging (MRI), spatial RNA sequencing (spRNAseq), functional BBB assessment, and immunofluorescence average-intensity-mapping. We identified a pronounced early response of the choroid plexus (CP) peaking at 12 to 24h, that was characterized by inflammatory cytokine expression, epithelial and endothelial expression of leukocyte adhesion molecules, and the accumulation of leukocytes. In contrast, we observed a delayed secondary reaction pattern at the injection site (striatum) peaking at 96h, defined by gene expression corresponding to perilesional leukocyte infiltration and correlating to the delayed signal alteration seen on MRI. Pathway analysis revealed a dependence of the early inflammatory reaction in the CP on toll-like receptor 4 (TLR4) signaling via myeloid differentiation factor 88 (MyD88). TLR4 and MyD88 knockout mice corroborated this observation, lacking the early upregulation of adhesion molecules and leukocyte infiltration within the CP 24h after whole-blood injection. In conclusion, we report a biphasic brain reaction pattern after ICH with a MyD88-TLR4-dependent early inflammatory response of the CP, preceding inflammation, edema and leukocyte infiltration at the lesion site. Pharmacological targeting of the early CP-activation might harbor the potential to modulate the development of ICH-SBI.

**Graphical abstract:** 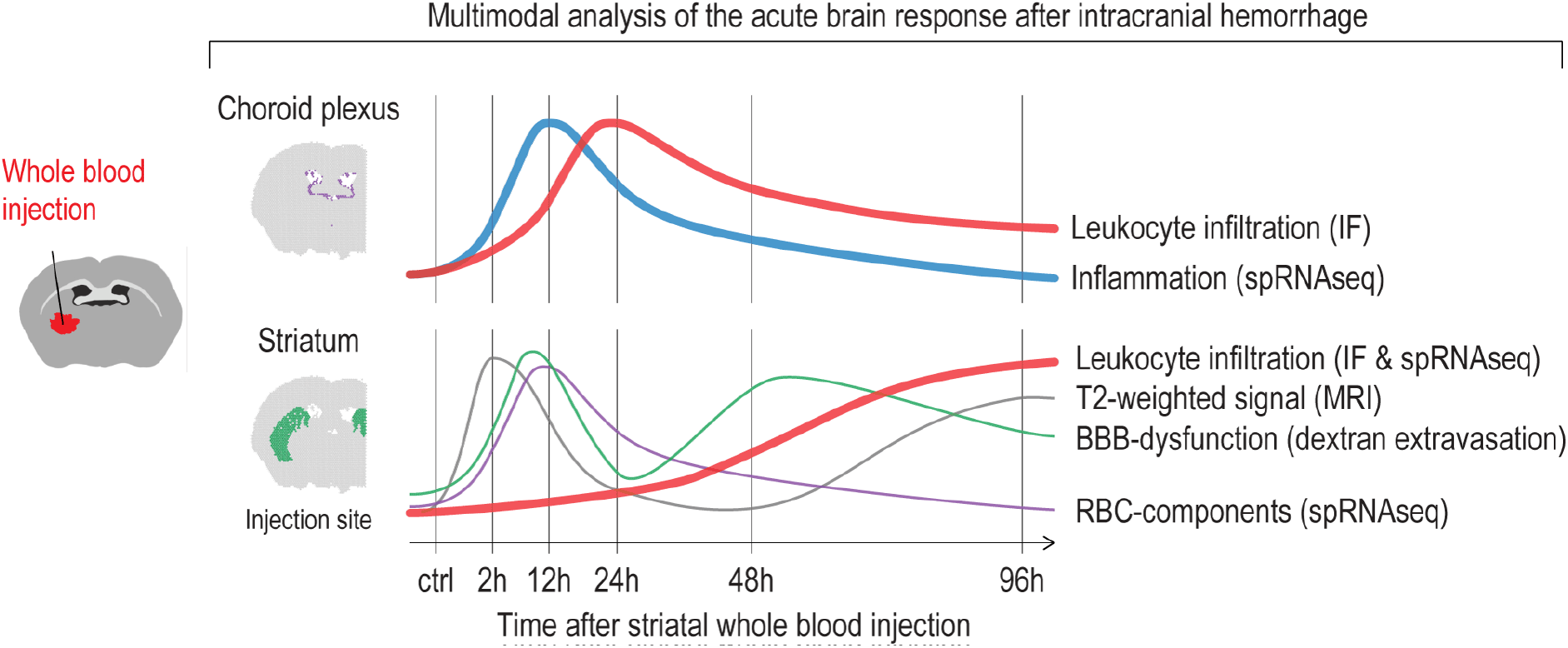

Akeret, Buzzi et al. characterized the spatiotemporal dynamics after striatal whole blood injection in mice using magnetic resonance imaging (MRI), spatial RNA sequencing (spRNAseq), functional blood-brain barrier (BBB) assessment, and immunofluorescence average intensity mapping (IF). They report a biphasic brain reaction pattern with an early MyD88-TLR4-dependent inflammatory response of the CP, which preceded secondary inflammation and leukocyte infiltration at the perilesional site.

## Introduction

Intracerebral hemorrhage (ICH) accounts for 15-20% of all strokes^1^. ICH is less common than ischemic stroke but results in substantially higher morbidity and mortality^2^. While the acute mass effect of a hemorrhage leads to immediate mechanical brain damage, functional neurological outcomes are strongly related to the occurrence of secondary brain injury (ICH-SBI) that evolves within days or a few weeks after the initial ICH^3^. Perihemorrhagic edema is a radiological surrogate for ICH-SBI that correlates with the need for surgical rescue measures and a poor clinical outcome^4^. Mechanisms that have been hypothesized to contribute to perihemorrhagic edema and ICH-SBI include the incitement of inflammatory pathways, dysfunction of the blood–brain-barrier (BBB), activation of resident microglia, and an influx of blood-borne immune cells^5^. However, the spatial and temporal interplay of specific inflammatory processes after ICH has not been sufficiently characterized, limiting targeted interventions to prevent and treat ICH-SBI^6^.

There are three compartments of decreasing immune privilege in the brain: brain parenchyma, cerebrospinal fluid (CSF) space, and leptomeninges, including the choroid plexus (CP) as an intraventricular evagination^7^. While the CP was long considered as an exclusively secretory organ for CSF, there is growing evidence for an immunological role in the pathogenesis of various neurological conditions^8^. The CP is a highly vascularized convolute located in the ventricular system composed of a monolayered epithelium and a stromal core^9^. Microvilli on the CSF side of the CP epithelium create an enormous apical surface area, which corresponds to approximately 10% of the BBB^9,10^. The perfusion of the CP stroma is the highest within the entire brain^11^. Given its central location in the brain, its enormous contact area with both the CSF and peripheral circulation, and its extensive perfusion, we hypothesize that the CP plays a pivotal role in the regulation of inflammatory processes after ICH.

Using a whole-blood striatal injection mouse model, this study explored the spatial and temporal dynamics of inflammatory processes after ICH using 7-Tesla magnetic resonance imaging (MRI), spatial RNA sequencing (spRNAseq), functional BBB assessment, and immunofluorescence average-intensity-mapping. Our data revealed a biphasic brain reaction pattern, including a distinct early inflammatory response of the CP, which preceded inflammation and leukocyte infiltration at the lesion site. The posthemorrhagic CP response was dependent on toll-like-receptor-4 (TLR4) signaling via myeloid-differentiation-factor-88 (MyD88).

## Methods

### Experiment series and animals

The experiments were organized into temporal and MyD88-TLR4 series. Temporal series investigated the spatiotemporal dynamics after striatal whole-blood injection using MRI, spRNAseq, and immunohistology. All temporal experiments were performed using 10-12-week-old wild-type C57BL/6J mice (*n_MRI_*=10, *n_spRNAseq_*=6, *n_Immunohistology_*=21). The inflammatory CP response findings determined the time point and specific readouts used in the MyD88-TLR4 series. For the latter, we used 10-12-week-old MyD88 (*n*=8) and TLR4 (*n*=6) knockout mice with wild-type littermate controls (*n_MyD88+/+_*=16, *n_TLR4+/+_*=12). All experiments were approved by the Swiss Federal Veterinary Office (ZH89/2019). Mice were obtained from Charles River Laboratories (Sulzfeld, Germany) and housed in individually ventilated cages within the Laboratory Animal Services Center at the University of Zurich with a 12-h/12-h dark/light cycle.

### Randomization and blinding

Group allocation (time points, injection fluid) was based on simple randomization using R (version 4.0.3)^12^. Investigators were blinded to group allocation.

### Striatal injection model

Whole-blood was collected from deeply anesthetized mice by terminal cardiac puncture, anticoagulated with sodium-citrate and used immediately. Stereotactic striatal needle insertion followed a previously published protocol^13,14^. We inserted a 33-G needle with a speed of 0.1 mm/s. 10 μL of fluid were injected with 2000 nL/min. The needle was left in place for 10 min, then removed slowly (0.1 mm/s) and the skin sutured. After surgery, the animals were placed in a heated wake-up box and monitored until they were fully recovered and could be transferred back to their cage.

### Magnetic resonance imaging

MRI data was collected at different time points (2h, 24h, 48h, 96 h) after surgery in the same mice, using a 7/16 small animal MR scanner (Pharmascan, Bruker Biospin GmbH, Ettlingen, Germany). Detailed technical and procedural information are provided in the Supplemental Material.

### Cryosections

For spRNAseq analysis, mice were transcardially perfused with 50 mL of ice-cold phosphate buffered saline (PBS) at different time points (2h, 12h, 24h, 48h, 96 h) after striatal whole-blood injection. The brain was extracted and immediately flash-frozen in cold Tissue-Tek OCT Compound (Sakura Finetek, Netherlands) using a liquid nitrogen-cooled metal block and stored at −80 °C until sectioning. We maintained the time from brain extraction to freezing below 1 min. Using a Leica CM3050 S Cryostat (Leica, Germany), 10-μm coronal cryosections were prepared and placed within the etched frames of the capture areas on the active surface of the Visium Spatial Slide (10X Genomics, Pleasanton, CA, USA).

### Spatial RNA sequencing

Tissue processing was performed according to the manufacturer’s instructions (User Guide CG000239, Rev A)^13^. Sections were fixed with cold methanol (−20 °C) and stained with H&E. Prior to permeabilization, we obtained brightfield images of the stained sections using an Axio Observer (Zeiss, Germany). Then, tissue sections were permeabilized for 6 min, followed by reverse transcription, second-strand synthesis, denaturation, complementary DNA (cDNA) amplification, and SPRIselect (Beckman Coulter, Brea, CA, USA) cDNA cleanup. Finally, cDNA libraries were prepared and sequenced on an Illumina NovaSeq 6000 (Illumina, San Diego, CA, USA) using a 28 bp + 120 bp paired-end sequencing mode.

Mapping and counting were performed using Space Ranger 1.1.0 (10X Genomics), with the reference genome GRCm38.p6 (mm10). Space Ranger aligned the barcodes in each read with a feature (i.e., a single spot with 55 μm diameter on the slide with a distinct spatial position defined by a unique barcode) relative to the fiducial frame, thereby associating read counts with spatial location and histological image. Downstream analysis was performed in python using the workflow and packages described in the Supplemental Material^15^.

### Immunohistology

Immunohistological analyses were coupled to an assessment of the BBB. Mice received an intraperitoneal injection of a fluorescently labeled dextran diluted in 0.9% saline solution (Supplemental Table 1) 2h before euthanasia. For euthanasia, mice were deeply anesthetized with intraperitoneal injection of ketamine/xylazine/acepromazine and transcardially perfused with 0.1M PBS followed by 30 mL ice-cold 4% paraformaldehyde in PBS. After removal of the brain and post-fixation overnight in 4% paraformaldehyde at 4 °C, brains were cut into 60-μm histological sections using a vibratome (VT1000 S; Leica, Geneva, Switzerland). For quantification, a standardized selection of eight sections across the lesion (every second section, beginning at 300 μm posterior to the anterior commissure) was collected. Incubation with primary antibodies was performed over 48h at 4 °C, followed by secondary antibodies over 24h at 4 °C and Hoechst staining for 40 min. Between each staining-step, slices were washed three times. After mounting with ProLong Gold (Invitrogen, CA, USA), imaging was performed using the Axio Scan.Z1 (Zeiss, Oberkochen, Germany). The antibodies and the dilutions used for staining are listed in Supplemental Table 2.

### Histological quantification

To achieve a reproducible representation of the lesion, the CP, and the cerebral hemisphere, histological quantification was performed on a selection of eight anatomically standardized coronal sections per mouse (every second section, beginning at 300 μm posterior to the anterior commissure). Leakage area size and intensity were semi-automatically quantified by two blinded investigators with a custom-made Fiji plugin^14^. Average-intensity-maps were calculated in a python environment after manual segmentation of the different anatomical compartments by two blinded investigators as described in the Supplemental Material.

### Data visualization and statistical analysis

Statistical analysis was performed using R (version 4.0.2)^12^. In the temporal series, for each time point a 95% bootstrap confidence interval was calculated and visualized as a box. As a summary, a loess curve was fitted to the data using a t-based approximation and plotted as a line. In the knockout series, the boxplots represent group averages on an animal level. The box bounds the interquartile range (IQR) divided by the median, while the whiskers extend to the highest and lowest value within the 1.5 IQR, respectively.

### Data and code availability

The Fiji-script, processed spRNAseq-data, python script, and R-code to reproduce all analyses and plots are available online (https://doi.org/10.5281/zenodo.5760872).

## Results

### Striatal whole-blood injection evokes a biphasic injury phenotype

First, MR-morphological and hematoxylin and eosin (H&E) histological changes were investigated 2h, 24h, 48h, and 96h after striatal whole blood injection (Figure 1A). Serial T2-weighted and susceptibility-weighted (SWI) images demonstrated an acute transient T2/SWI signal change at 24h (Figure 1B). A secondary phase of T2/SWI-hypointensity was consistently detected at 96h. The H&E appearance of the lesion remained largely unchanged up to 48h post-injection with only sparse cellular infiltration (Figure 1C). From 48h to 96h, there was a substantial reduction of the blood clot area associated with increased cellularity at the lesion site. Collectively, MRI demonstrated a biphasic injury phenotype with a secondary signal alteration coinciding with histological evidence of cellular infiltration and hematoma resolution.

**Fig 1.**
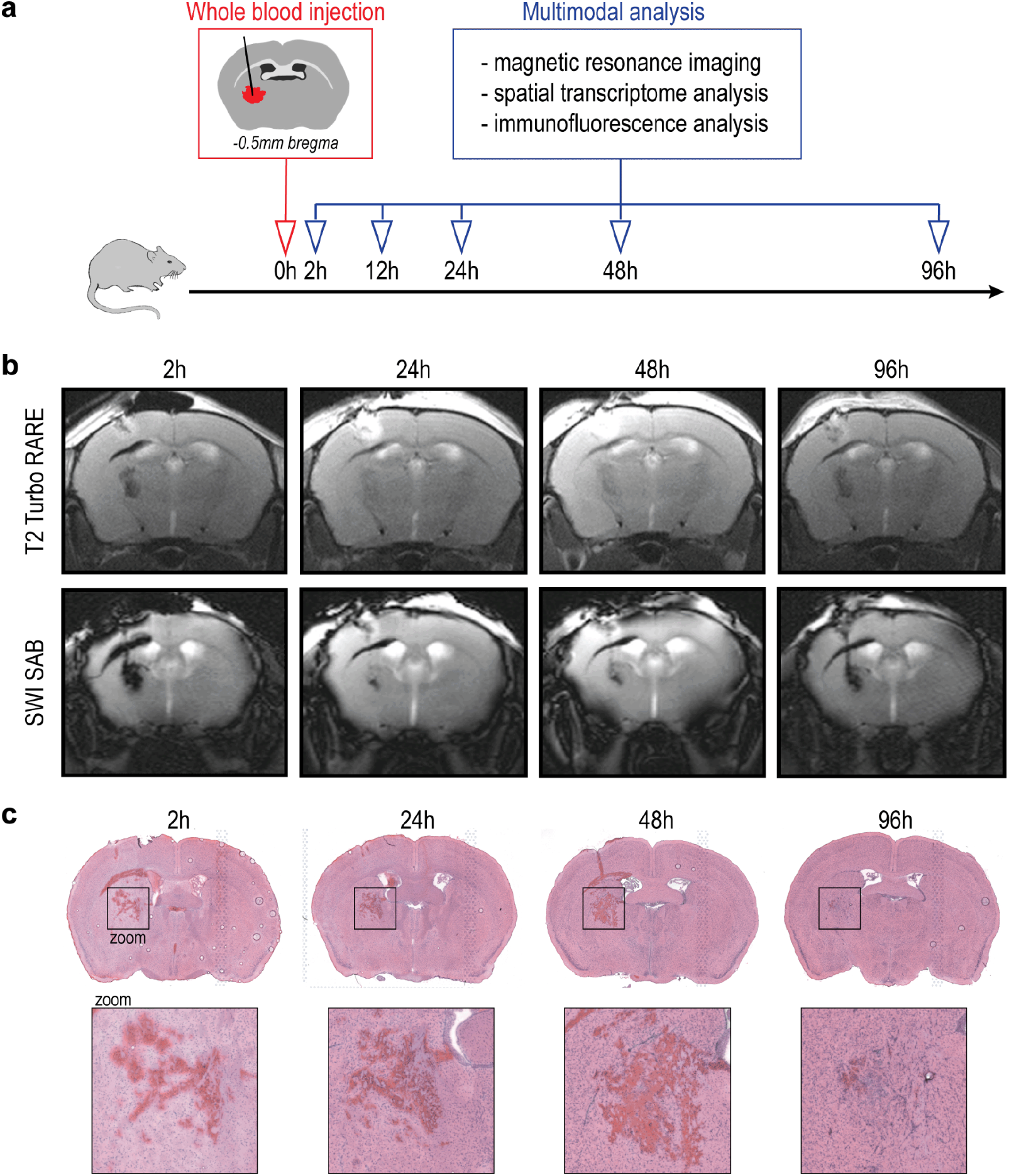
MRI and histological changes after striatal whole-blood injection. **A.** Schematic representation of the study design. **B.** T2-weighted and susceptibility-weighted (SW) MR images 2h, 24h, 48h, and 96h after striatal whole-blood injection (representative example from *n*=10). **C.** Lesion site over time on H&E-stained coronal cryosections.

### Transcriptional changes to striatal whole-blood injection

Next, we explored spatial transcriptome changes after striatal whole-blood injection applying the 10x Genomics workflow while focusing on the brain regions comprising the perilesional area (striatum) and the CP. This approach allowed the extraction of gene expression data from these specific brain regions, which are otherwise difficult to selectively isolate. We obtained spRNAseq data from a control mouse (no injection) and from mice 2h, 12h, 48h, and 96h after whole-blood injection. Per-spot gene expression data from all animals were merged into a combined dataset comprising 22’835 features. Each feature represented the gene expression pattern of the underlying tissue with a diameter of 55 μm. Aiming at an unbiased spatial segmentation of the striatum and CP, we used unsupervised clustering and predefined gene sets^16^ to assign each feature to an anatomical region. Figure 2A illustrates an Uniform Manifold Approximation and Projection (UMAP) with a Leiden clustering restricted to five main clusters. We calculated gene scores for eight different anatomical regions (Figure 2B). Four of the five clusters (1-4) matched to a single region (Figure 2C). Cluster 0 (blue) had an association with four regions (pallidum, hypothalamus, midbrain, pons), collectively defined as deep brain structures. The correct anatomical annotation of the gene expression features was confirmed by back-projection and anatomical data visualization (Figure 2D, Supplemental Figure 2A).

**Fig 2.**
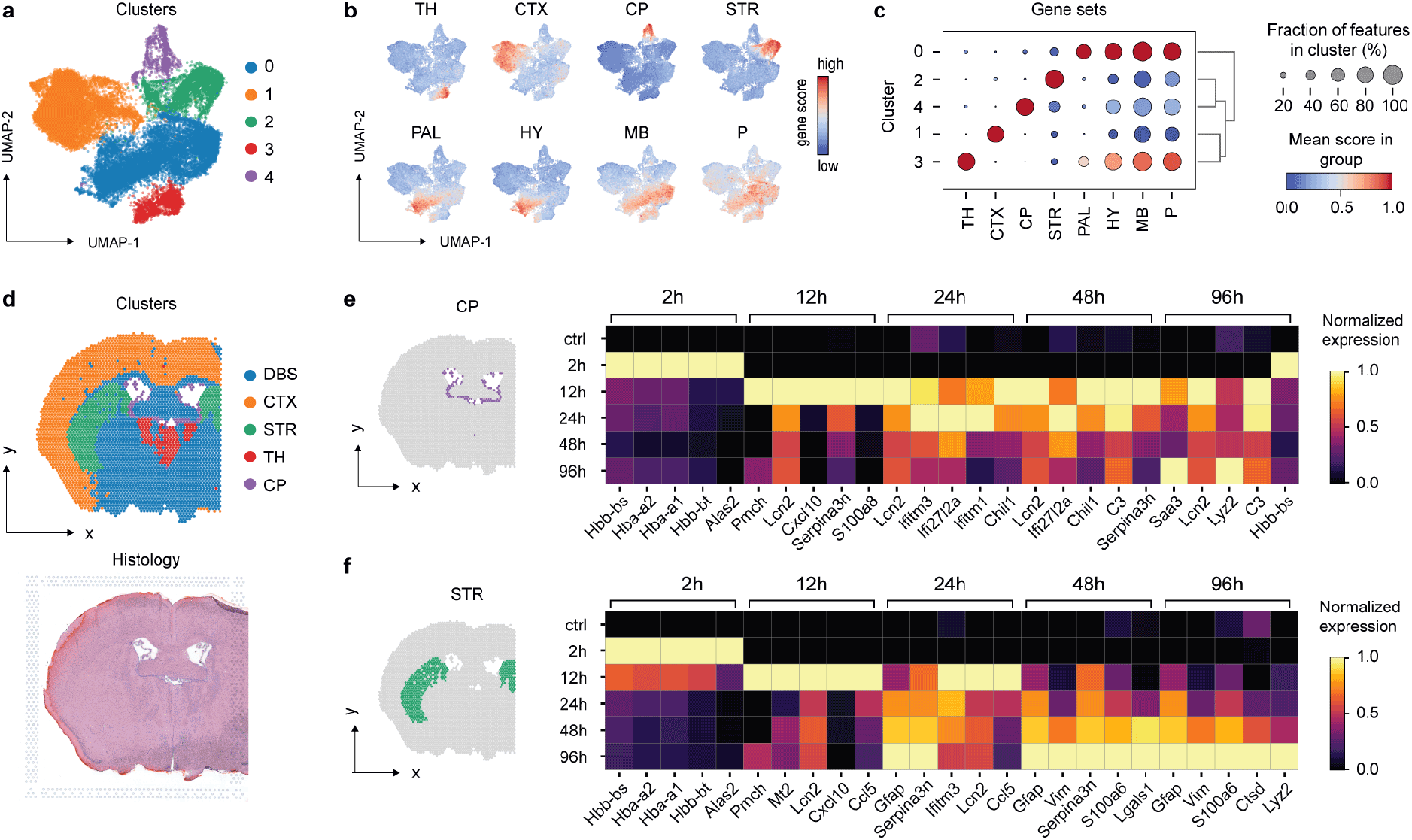
Spatiotemporal differential gene expression after striatal whole-blood injection. **A.** UMAP plot colored by the unsupervised clustering of the merged spatial transcriptome dataset, including all features from *n*=6 mice (no striatal injection [control], different time points after striatal whole-blood injection [2h, 12h, 24h, 48h, 96 h]). **B.** Gene set scores for specific anatomical compartments projected on the UMAP plot. *TH, thalamus; CTX, cortex; CP, choroid plexus; STR, striatum; PAL, pallidum; HY, hypothalamus; MB, midbrain; P, Pons.* **C.** Mean anatomical gene set expression score (*dot color*) and fraction of positive features (*dot size*) of each identified cluster. **D.** Spatial projection of the identified clusters on the control sample (top) and corresponding H&E histology (bottom). **E.** Top five differentially expressed genes (control as baseline) in the CP cluster stratified by time point after injection. **F.** Top five differentially expressed genes (control as baseline) in the STR stratified by time point after injection.

Objective anatomical segmentation enabled a regionally stratified analysis of temporal gene expression changes. Within the five regions, we performed a differential gene expression analysis comparing each post-injection time point with the untouched control mouse (Figure 2E&F, Supplemental Figure 2B). Heatmaps in Figures 2E and 2F illustrate the expression of the top five differentially expressed genes per time point and region. The CP and the striatum shared a very similar early transcriptional signal, with gene expressions of red blood cell components (hemoglobin, Alas2) peaking at 2h post-injection, followed by a number of inflammatory gene expressions spiking at 12h. This indicated that both regions were directly exposed to the injected blood or its degradation products and that these exposures evoked an acute transcriptional reaction. The red blood cell signal disappeared within 24h, coinciding with the disappearance of the immediate post-injection signal alteration in MRI. In contrast to this early phase, the CP and the striatum differed in their gene expression patterns at later time points. The CP demonstrated a monophasic activation: After a clearly defined first gene expression wave spiking at 12-24h, only few genes were reaching an expression peak at 48 or 96h. In contrast, the striatum demonstrated a biphasic response, with a defined secondary wave of gene expression peaking at 96 h. This delayed response correlated with the second signal alteration on MRI at 96h and is consistent with a delayed inflammatory reaction at the injection site.

### Early inflammatory signaling in the CP following striatal whole-blood injection

Using hallmark genes, we investigated the principal biological processes contributing to the transcriptional responses in the CP and the perilesional site. The marker genes, identified by the differential gene expression analysis, could be assigned to six different biological processes (Figure 3A). The initial cytokine upregulation (e.g., Cxcl10, Ccl5), with a peak at 12h, was similar in both compartments. However, compared to the perilesional area, the CP featured a more pronounced early increase of interferon response elements (e.g., Irf7, Ifi27l2a) (Figure 3B) and cell adhesion molecules (e.g., Vcam1, Bst2) (Figure 3C) peaking between 12 and 24h. This was accompanied by a peak in the inflammatory response genes (Serpina3n, Lcn2). The perilesional site, in contrast, displayed a delayed increase in the same inflammatory markers peaking at 96h (Figure 3A). This delayed response of the perilesional site was accompanied by an increased expression of macrophage markers (Cd68, Lyz2) (Figure 3D) and genes involved in heme/iron metabolism (Hmox1, Ftl1). The early inflammatory reaction with cytokine signaling, adhesion molecule expression, and leukocyte attraction in the CP was confirmed by selected marker genes. Both Ccl20, a chemokine expressed by the CP epithelium but not the brain parenchyma^17^, and the prototypical leukocyte adhesion molecule, Icam1, demonstrated a peak in the CP 12h after striatal whole-blood injection (Figure 3E&F). The pan-leukocyte marker Ptprc (Cd45) showed a biphasic CP expression with an initial increase at 24h followed by a second peak at 96h (Figure 3G). Spatial gene expression revealed an acute inflammatory CP activation characterized by upregulation of interferon response elements, CCL20, adhesion molecules, and leukocyte markers. This acute CP response precedes the macrophage-dominated inflammation in the perilesional area.

**Fig 3.**
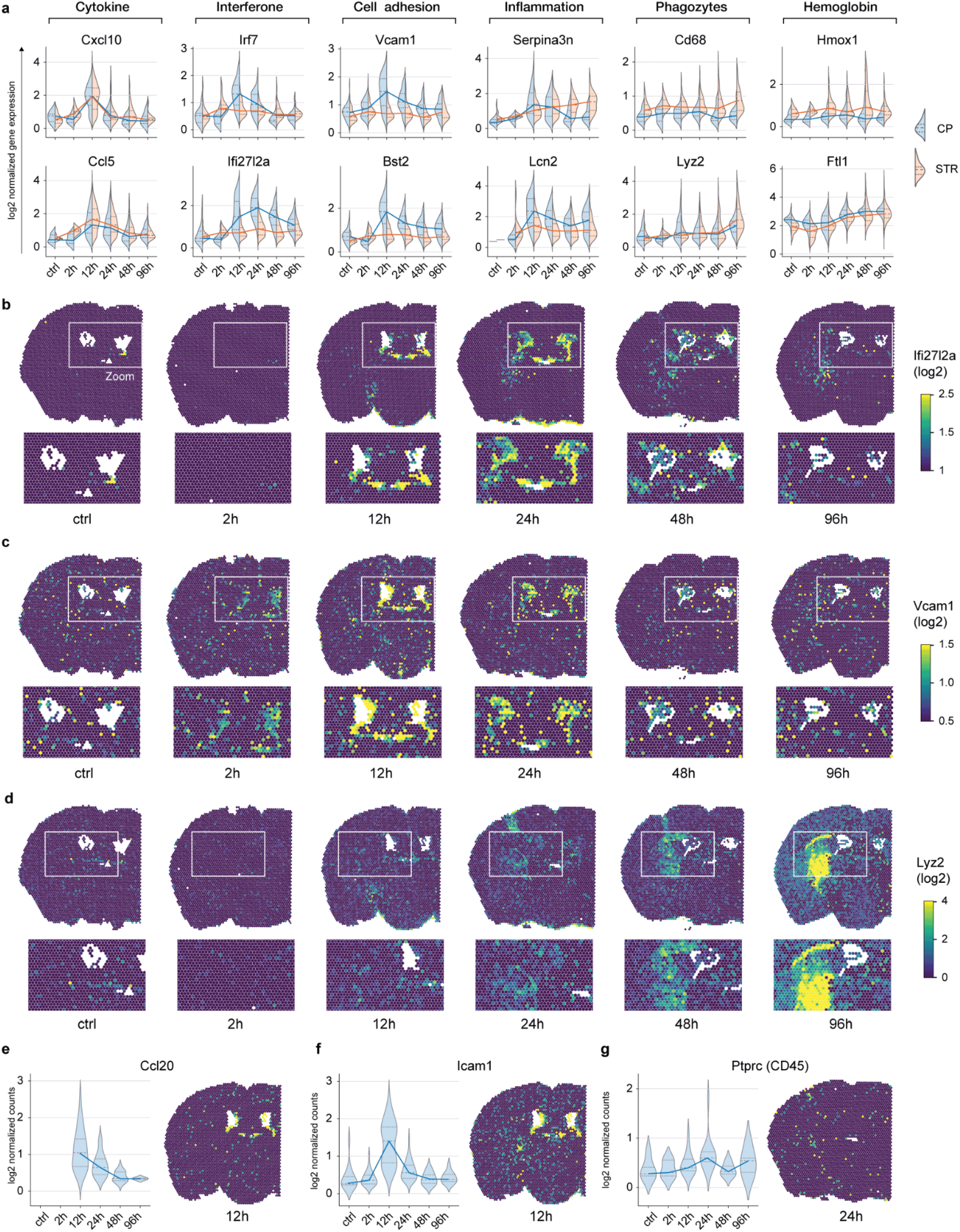
Perilesional and CP transcription profiles after striatal whole-blood injection. **A.** Violin plots of the temporally resolved log2-normalized expression of selected genes in the CP (violet) and the striatum (STR) (green) (*n*=6). **B.-D.** Spatial projection of the log2 normalized expression of Ifi27l2a (**B**), Vcam1 (**C**), and Lyz2 (**D**) in the control sample and at different time points after whole-blood injection. **E-G.** Temporal dynamics of the log2 normalized expression of Icam1 (**E**), Ccl20 (**F**) and Ptprc (CD45) (**G**) in the CP visualized as violin plots and spatial gene expression plot.

### Inflammatory CP activation precedes secondary perilesional blood–brain barrier disruption and leukocyte infiltration

Next, we quantified BBB function, leukocyte infiltration, and markers of parenchymal inflammation through the generation of average-intensity-maps 2h, 12h, 24h, 48h, and 96h after striatal whole-blood injection, representing a quantitative summary of fluorophore accumulation and distribution (Figure 4A). The permeability of the BBB was measured by the perilesional extravasation of dextran (10 kDa, 70 kDa, and 155 kDa). The extravasation area size (Figure 4B&C) and the mean fluorescence intensity (Supplemental Figure 2A) showed a biphasic temporal course. The extravasation area size had a first peak after 12h and a second peak after 48 h. The extravasation mean fluorescence intensity peaked 2 and 48h after injection. The fluorescent signal of dextran in the CP differed between anatomical locations with the highest values in the ipsilateral CP (CP_ipsi_>CP_third_>CP_contra_) at the 2h time point (Supplemental Figure 2B-D). The biphasic temporal profile of dextran extravasation is best explained by an acute mechanical disruption of the perilesional BBB causing the first leakage peak, followed by an inflammation-driven secondary spike around 48h after injection.

**Fig. 4.**
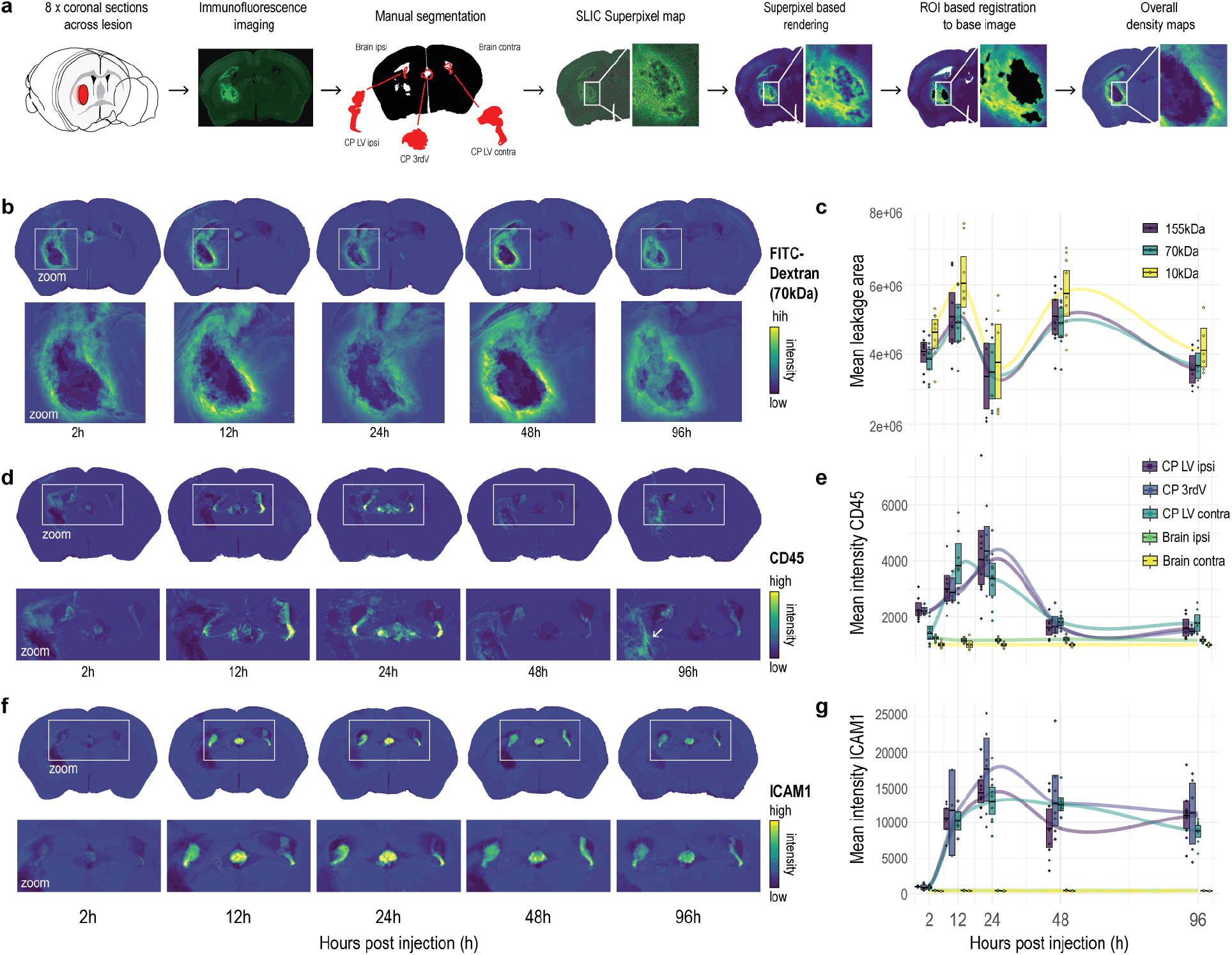
Dextran extravasation and inflammatory protein expression after striatal whole-blood injection. **A.** Methodological steps of the semi-automated quantification of spatial protein expression (*n*=21). **B.** FITC-Dextran (70 kDa) average-intensity-maps at different time points with perilesional zoom-in. **C.** Temporal dynamics of the mean leakage area for different dextrans (155 kDa, 70 kDa, 10 kDa). **D.** CD45 average-intensity-maps at different time points with a CP/perilesional zoom-in. The arrow at 96h indicates the high perilesional CD45 signal intensity. **E.** Temporal dynamics of the mean CD45 signal intensity in different anatomical compartments. *LV, lateral ventricle; 3rdV, third ventricle.* **F.** ICAM1 average-intensity-maps at different time points with CP zoom-in. **G.** Temporal dynamics of the mean ICAM1 signal intensity in different anatomical compartments.

Leukocyte infiltration and parenchymal inflammation were visualized by immunofluorescence for the pan-leukocyte antigen CD45 and the adhesion molecule ICAM1. Compared to all other brain regions, there was a uniquely intense CD45 signal in the CP peaking at 24h (Figure 4D&E). In contrast, the CD45 signal at the site of the hematoma displayed a delayed appearance starting at 48h and markedly increasing at 96h after injection (Figure 4D, arrow), coinciding with the secondary lesion phenotype changes seen on MRI and in RNA expression. Similarly, ICAM1 displayed an intense and acute expression in the CP but not in the other brain regions. A strong ICAM1 signal was already clearly detectable after 12h and reached its peak at 36h post-injection (Figure 4F&G, Supplemental Figure 2E shows co-staining for collagen IV as internal control).

Representative topographic antigen representation by confocal microscopy is given in Figure 5. We found scarce CD45-positive cells within the CP 2h post-injection, while CD45-positive cells were abundant in the CP stroma at 24h (Figure 5A). These cells were positive for F4/80 but negative for Iba1 and TMEM119, consistent with a phenotype of blood monocyte-derived macrophages (Figure 5B). Correspondingly, ICAM1 expression was low in the CP 2h after whole-blood injection, followed by a striking increase after 24h. ICAM1 was primarily expressed on the CSF surface of the CP epithelium (Figure 5C). At 24h post-injection, abundant Iba1-positive, F4/80-negative, and TMEM119-negative cells appeared at the CSF surface of the CP epithelium (Kolmer’s epiplexus cells) (Figure 5D). Coincident with the macrophage infiltration signal in the spRNAseq data and the delayed signal alteration in MRI, we observed a perilesional increase of CD45-positive cells at 96h (Figure 5E).

**Fig 5.**
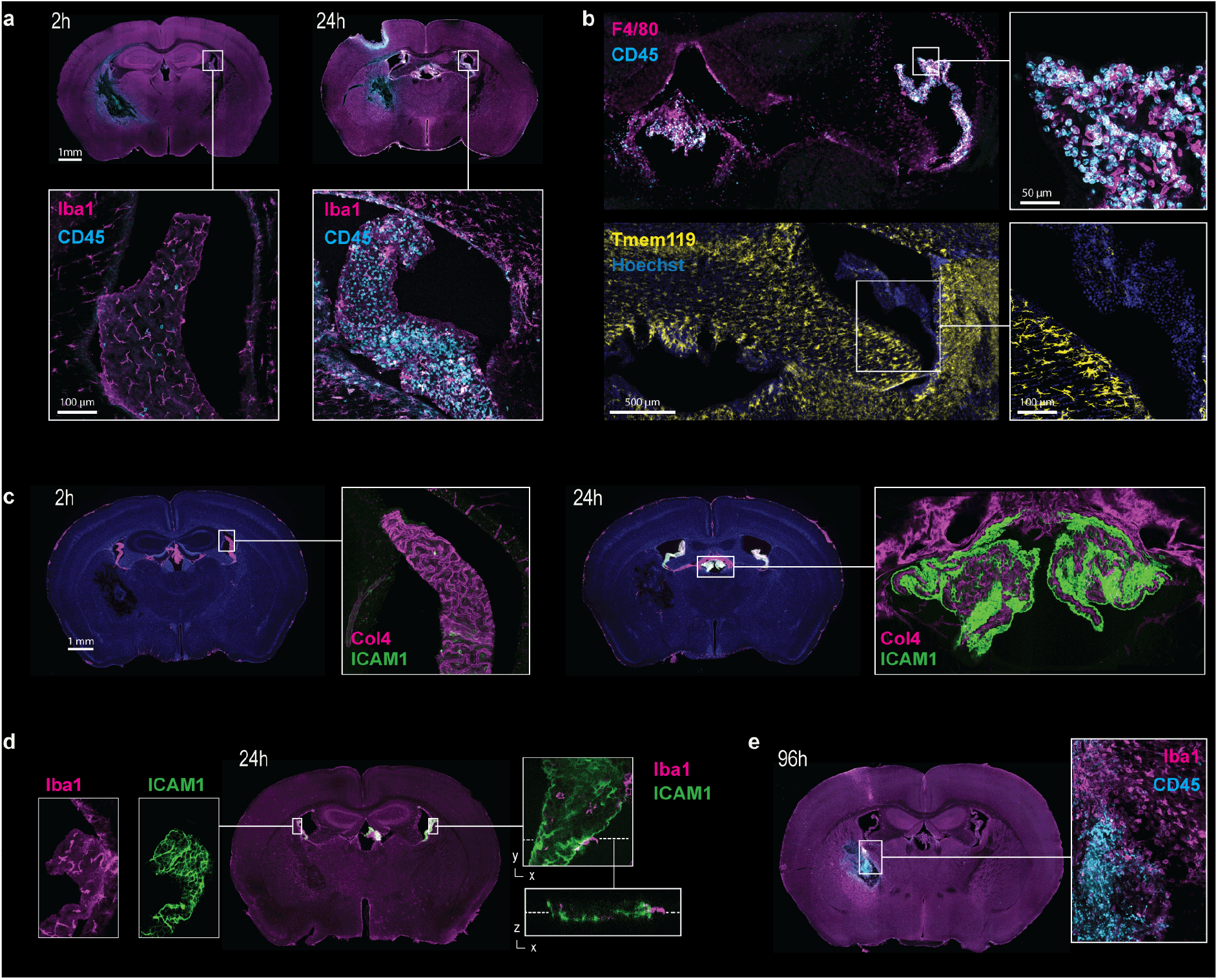
Representative immunohistology after striatal whole-blood injection. Representative vibratome sections at various time points after whole-blood injection. **A.** 2h (left) and 24h (right) stained for Iba1 (magenta) and CD45 (cyan). **B.** 24h stained for F4/80 (magenta) and CD45 (cyan) or TMEM119 (yellow) and nuclei (blue). **C.** 2h (left) and 24h (right) stained for collagen IV (Col4, magenta) and ICAM1 (cyan). **D.** 24h stained for Iba1 (magenta) and ICAM1 (green). **E.** 96h stained for Iba1 (magenta) and CD45 (cyan).

Collectively, immunohistological analyses showed that striatal whole-blood injection triggered an acute inflammatory response of the CP with pronounced epithelial ICAM1 expression, epiplexus macrophage lining, and stromal leukocyte invasion. This acute CP response was followed by delayed secondary perilesional dextran extravasation and leukocyte infiltration.

### MyD88-TLR4 gene deletion attenuates acute inflammatory CP activation after striatal whole-blood injection

Finally, we defined the signaling pathway linking striatal whole-blood injection and the inflammatory activation of the CP. Enrichment analysis of the spatial gene expression data set, using selected *KEGG 2019 mouse^18^* gene sets, identified significant enrichment of the toll-like receptor signaling pathway (Figure 6A). More specifically, we found a distinct temporal association of the acute inflammatory CP response with the post-injection dynamics of the scored expression of a MyD88-dependent TLR signaling gene set (GO:0002755)^19^ (Figure 6B&C). This analysis was consistent with the outcome of experiments in TLR4 and MyD88 knockout mice (Figure 6D-G). Compared to wild-type littermates, ICAM1 expression and infiltration of F4/80-positive leukocytes at 24h post whole-blood injection were markedly reduced in the CP of both mouse models, almost to the level observed in sham animals injected with saline.

**Fig 6.**
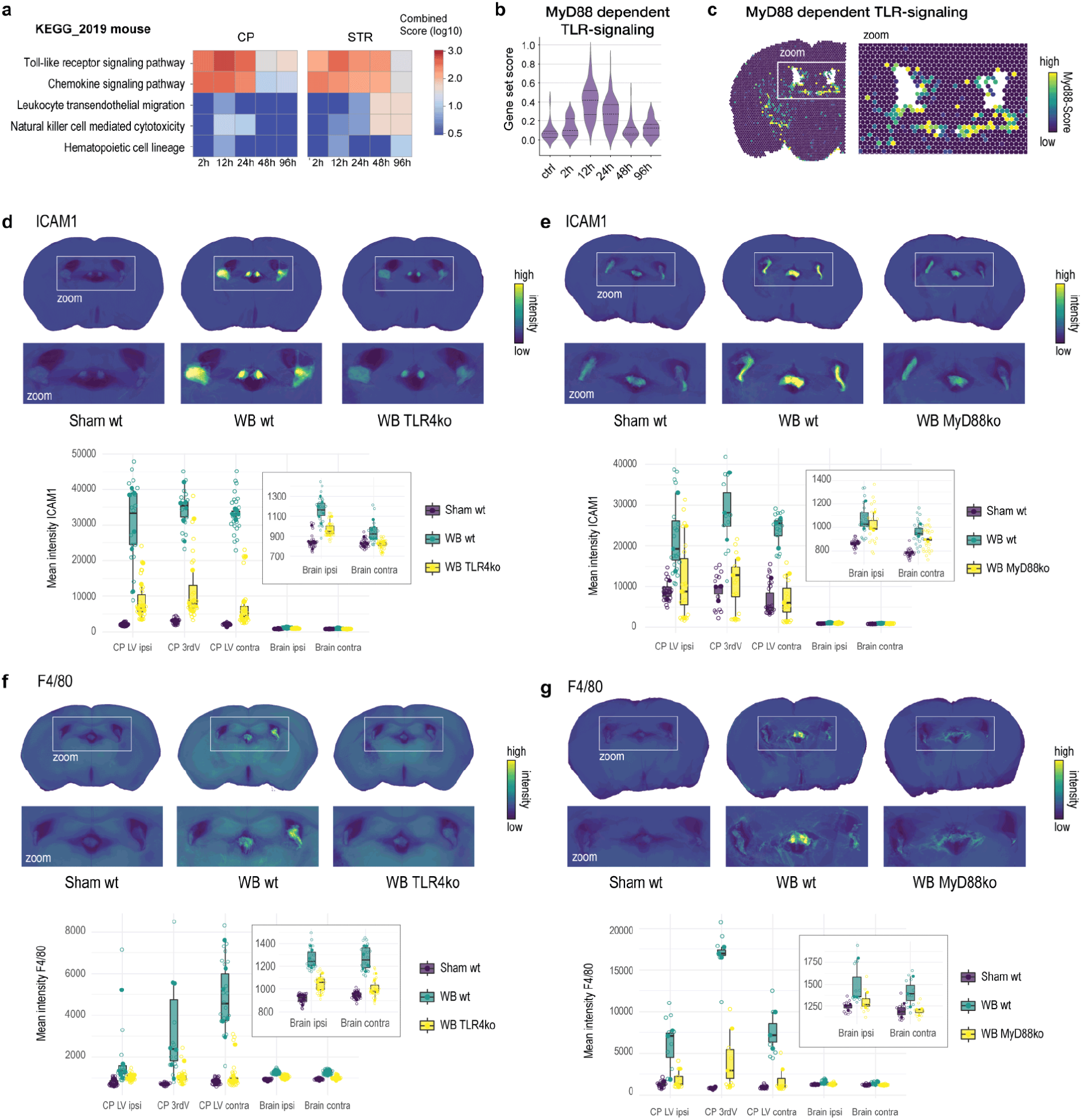
MyD88-TLR4 dependence of the CP response after striatal whole-blood injection. **A.** *KEGG-2019* mouse-based gene-set enrichment patterns in the CP and the striatum (STR) (control as baseline, *n*=6). **B.** Temporal dynamics of the *MyD88-dependent-TLR4-signaling* gene set score in the CP (*n*=6). **C.** Spatial projection of the *MyD88-dependent-TLR4-signaling* gene set score 24h after injection. **D.** ICAM1 signal intensity 24h after striatal NaCl (sham, *n*=6) or whole-blood (*n*=6) injection in littermate TLR4+/+ mice (wt), and whole-blood injection in TLR4-/- (TLR4ko) mice (*n*=6). Top: ICAM1 density-maps with a CP zoom-in. Bottom: Mean ICAM1 signal intensity in different anatomical compartments. *LV, lateral ventricle; 3rdV, third ventricle.* **E.** ICAM1 signal intensity 24h after striatal NaCl (sham) or whole-blood injection in littermate MyD88+/+ mice (wild-type), and whole-blood injection in MyD88 knockout (MyD88ko) mice (*n*=8 per group). Top: ICAM1 density-maps with a CP zoom-in. Bottom: Mean ICAM1 signal intensity in different anatomical compartments. **F.** F4/80 signal intensity 24h after striatal NaCl (sham) or whole-blood injection in littermate TLR4+/+ mice (wild-type), and whole-blood injection in TLR4 knockout (TLR4ko) mice (*n*=6 per group). Top: F4/80 density-maps with a CP zoom-in. Bottom: Mean F4/80 signal intensity in different anatomical compartments. **G.** F4/80 signal intensity 24h after striatal NaCl (sham or whole-blood injection in littermate MyD88+/+ mice (wt), and whole-blood injection in MyD88-/- (MyD88ko) mice (*n*=8 per group). Top: F4/80 density-maps with a CP zoom-in. Bottom: Mean F4/80 signal intensity in different anatomical compartments.

## Discussion

This study explored the spatial and temporal dynamics of inflammatory processes after ICH in a striatal whole-blood injection mouse model using 7-Tesla MRI, spRNAseq, functional BBB assessment, and immunofluorescence average-intensity-mapping. Consistent with the delayed onset of perihemorrhagic edema in ICH patients, we observed a delayed secondary perilesional inflammation and BBB dysfunction in our model. On MRI, this was evidenced by changes in the T2-weighted signal intensity at the injection site after 48-96h. This imaging pattern correlated with a delayed increase in perilesional dextran extravasation, upregulation of proinflammatory cytokine genes, recurrent astrocyte activation, and pronounced macrophage infiltration.

Secondary perilesional inflammation was preceded by a distinct inflammatory activation of the CP peaking at 12-24h post-injection. We identified a remarkable upregulation of leukocyte adhesion molecules on the apical surface of the CP and an increase in the number of Kolmer’s epiplexus cells within hours after striatal whole-blood injection. Simultaneously, there was a marked stromal accumulation of F4/80-positive and TMEM119-negative leukocytes, presumably reflecting the peripheral blood recruitment of monocyte-derived macrophages. CP epithelial cells reportedly possess the capacity to produce chemokines for systemic immune cell attraction upon both central and peripheral noxious stimuli^20^. Our spRNAseq data showed an early and high CP expression of chemoattractants, most dominantly CXCL10 and CCL5. CXCL10 and CCL5 attract blood monocytes and lymphocytes and promote their endothelial adhesion^21,22^. Given its strategic anatomical location, its enormous contact area with both the CSF and peripheral circulation, and its extensive perfusion, the CP might fulfill a pivotal function in CNS immune surveillance and serve as an interface between peripheral leukocytes and inflammation in the brain, possibly promoting secondary leukocyte infiltration of the lesion site^9^.

We established that the pronounced inflammatory activation of the CP after striatal whole-blood injection is dependent on MyD88-TLR4 signaling. There is a high level of TLR4 expression on the CP epithelium^23^. In addition, CSF hypersecretion after intraventricular whole-blood injection depends on TLR4 signaling in the CP epithelium^24^. Our spRNAseq data demonstrated a distinct temporal association between MyD88-dependent TLR signaling in the CP and its expression of adhesion molecules and chemoattractants. In MyD88 and TLR4 knockout mice, there was neither CP surface activation nor stromal leukocyte invasion after striatal whole-blood injection. In line with our findings, ICAM1, CCL20, CXCL10, and CCL5 expressions reportedly depend on MyD88-TLR4 signaling in other organs^25–28^. The CP thus appears to respond in a TLR4-dependent manner to intracranial hemorrhage with epithelial upregulation of adhesion molecules, release of centrally and peripherally oriented chemoattractants, and CSF hypersecretion.

Future studies are needed to define the nature of the specific TLR4 ligands and the precise site of their action. Currently, we can speculate that the tissue damage caused by the acute bleeding releases multiple tissue damage-associated molecular patterns (DAMPs) into the interstitial space of the brain. Additionally, the lysis of red blood cells may liberate large quantities of cell-free hemoglobin, which is prone to auto-oxidation and release of the TLR4 agonist heme^14,29–31^. We detected high levels of hemoglobin subunit transcripts in the CP as early as 2h after the striatal injection of whole-blood. If taken as a surrogate marker for all locally released DAMPs, this observation suggests a rapid transfer of signaling molecules from the lesion site to the CP epithelium, which acts in this mechanistic model as a sensor for remote brain injury, triggering downstream inflammation, CSF hypersecretion, and leukocyte infiltration from the peripheral circulation.

There are certain limitations to be considered while interpreting the results of our study. We based our data solely on a whole-blood injection model for ICH. Injection of bacterial collagenase is another established ICH model in rodents^32^. We adopted the whole-blood injection model because it allowed a more isolated investigation of brain exposure to blood, the relative hematoma volume could be easily adjusted to the human clinical correlate, and the lesion size demonstrated high reproducibility. In addition, although the blood was injected under stereotactic guidance into the right striatum, blood bursting into the ventricle in certain cases cannot be ruled out. While this results in some variability of the model, it is consistent with the clinical situation in humans, where 40% of patients with ICH have intraventricular blood^33^.

Collectively, this study enhances our understanding of the spatial and temporal dynamics of inflammatory processes after ICH. The pronounced and early reaction of the CP, preceding secondary perilesional inflammation and edema, positions the MyD88-TLR4-dependent immunological CP activation as a potential target to modulate ICH-SBI.

## Non-standard Abbreviations and Acronyms

BBB: blood–brain-barrier
CP: choroid plexus
ICH-SBI: intracerebral hemorrhage related secondary brain injury
MRI: magnetic resonance imaging
MyD88: myeloid differentiation factor 88
PBS: Phosphate buffered saline
ROI: region of interest
spRNAseq: spatial RNA sequencing
TLR4: Toll-like receptor 4.

## Acknowledgments

We thank the staff of the Functional Genomic Center Zurich for their support with transcriptome analysis, Marc-Aurel Augath of the Institute for Biomedical Engineering (University of Zurich and ETH Zurich) for his technical assistance during the MRI studies.

## Author contributions

*Conceptualization*: KA, RMB, MH, DJS; *Methodology*: KA, RMB, JK; *Investigation*: KA, RMB, BRT, NiS, NaS, LB, KH; *Formal analysis:* KA, RMB, BRT; *Visualization and Writing original draft*: KA, RMB, *Writing - review and editing*: all authors; *Funding acquisition*: KA, RMB, FV, MH, DJS; *Resources*: JK, MH, DJS.

