## Supplemental Material for "MyD88-TLR4-dependent choroid plexus activation precedes perilesional inflammation and edema in intracerebral hemorrhage"

* MH and DJS share last authorship

**Affiliations**

1 Department of Neurosurgery, Clinical Neuroscience Center, Universitätsspital and University of Zurich, Zurich, Switzerland

2 Division of Internal Medicine, Universitätsspital and University of Zurich, Zurich, Switzerland

3 Institute for Biomedical Engineering, University of Zurich and ETH Zurich, Zurich, Switzerland

**Short title**

Choroid plexus activation precedes ICH-SBI

**Corresponding author**

Michael Hugelshofer

Frauenklinikstrasse 10, 8091 Zürich, Switzerland

### Supplemental Methods

#### Magnetic resonance imaging

MRI data was collected at different time points (2 h, 24 h, 48 h, 96 h) after surgery in the same mice, using a 7/16 small animal MR scanner (Pharmascan, Bruker Biospin GmbH, Ettlingen, Germany) equipped with an actively shielded gradient set of 760 mT/m with an 80 µs rise time and operated using the Paravision 6.0 software platform (Bruker Biospin GmbH). A circular polarized volume resonator was used for signal transmission, and an actively decoupled mouse brain quadrature surface coil with integrated combiner and preamplifier was used for signal reception (Bruker BioSpin GmbH). Mice were anesthetized with an initial dose of 4% isoflurane (Abbott, Cham, Switzerland) in oxygen/air (200:800 ml/min) mixture and anesthesia was maintained with 1.5% isoflurane in oxygen/air mixture (100:400 ml/min), administered via a nose cone. Mice were placed in the prone position in the MR scanner. Body temperature was monitored with a rectal probe and kept within 36.5 ± 0.5 °C. Anatomical images were obtained for geometrical planning. T2-weighted images were acquired with a Turbo RARE spin-echo sequence. Twenty contiguous axial slices were acquired with a slice thickness of 0.7 mm; field-of-view of 20 mm × 20 mm; image size of 256 × 256, to achieve a nominal; and a spatial resolution of 78 µm × 78 µm. Data were acquired with an echo time of 33 ms, a relaxation time of 2200 ms, a RARE factor of 8, and three averages, within an acquisition time of 3 min 31s. For SWI, a two-dimensional flow compensated gradient-recalled echo (FLASH) sequence was applied with the following parameters: 12 slices, slice thickness of 0.8 mm, interslice distance 1.3 mm, field-of-view of 20 × 20 mm; image size of 256 × 256, resulting in a resolution of 78 µm × 78 µm. One echo with an echo time of 12 ms; repetition time of 250 ms; flip angle of 15°; and the number of averages of 15, within an acquisition time of 11 min 40 s was recorded. Field map-based shimming was performed prior to data acquisition using the automated MAPshim routine to improve the homogeneity of the magnetic field. SWI were computed as described previously.[1] The SWI processing module in ParaVision 6.0.1 (Bruker, Ettlingen, Germany) was used with a Gauss broadening of 1 mm and a mask weighting of 4.

#### Downstream spRNAseq analysis

For downstream processing of the spRNAsew data, we used the Scanpy python package (version 1.7.0)[2] following a standard processing pipeline for visualization data processing and downstream analysis. Data were normalized using a scaling factor of 1,000, log-transformed, variable genes with "Seurat flavor" were detected, principal components analysis was performed, a neighborhood graph was built, and the UMAP was calculated. Clusters were defined using the Leiden algorithm. We used the Scanorama package (version 1.7.1) to merge different datasets.[3] Differentially expressed genes in each comparison were defined using the Scanpy function "rank_genes_clusters" (Wilcoxon's rank-sum test; cut offs: log fold change > 2, adjusted p-value < 0.01). Pathway and gene set enrichment analyses were performed for these genes using the GSEApy (version 0.10.7) package.[4]

#### Histological quantification

Histological quantification was performed on a selection of eight anatomically standardized coronal sections per mouse (every second section, beginning at 300 µm posterior to the anterior commissure) to achieve a reproducible representation of the lesion, the CP, and the cerebral hemisphere. Leakage area size and intensity were semi-automatically quantified by two blinded investigators with a custom-made plugin[5] for the Fiji system.[6] Using the image processing software QuPath (version 0.2.3),[7] two blinded investigators manually segmented each histological slice into five anatomical regions of interest (ROI) relative to the lesion site (CP of the ipsilateral lateral ventricle ($CP_{ipsi}$), CP of the third ventricle ($CP_{third}$), CP of the contralateral lateral ventricle ($CP_{contra}$), ipsilateral brain parenchyma ($H_{ipsi}$), and the contralateral brain parenchyma ($H_{contra}$)). The mean signal intensity was determined (preferred pixel size of 1 µm) for each ROI and channel. Finally, group average intensity maps were generated for every channel; a whole-slice SLIC superpixel map (Gaussian sigma 5 µm, superpixel spacing 50 µm, 10 iterations, regularization 0.25) was generated. We calculated the mean channel intensity per superpixel (preferred pixel size of 0.1 µm), and the whole-slice measurement map was rendered as an RGB (red, green, and blue) image. One image per series and channel was selected as the reference image. Based on the manual segmentations, all ROIs were individually registered to the corresponding ROI of the reference image. Ultimately, group average ROIs were generated, which were fused based on maximal pixel intensity.

#### Supplemental Tables

**Supplemental Table 1. Details of the different fluorescently labeled dextrans used to assess the blood–brain barrier function.**

The specific antibodies used for immunostaining and their dilutions. Doses are given in µg per g bodyweight. *Tetramethylrhodaminisothiocyanat, TRITC; fluorescein isothiocyanate, FITC; Alexa Fluor 647, AF647.*

| Dextran (size) | Fluorphore | Dose | Company (Catalog #) |
|---|---|---|---|
| 155 kDa | TRITC | 125 µg/g | Sigma Aldrich (T1287-100M) |
| 70 kDa | FITC | 125 µg/g | Sigma Aldrich (FD70S-100MG) |
| 10 kDa | AF647 | 2.5 µg/g | ThermoFisher (D22914) |

**Supplemental Table 2. The specific antibodies used for immunostaining and their dilutions.**

| Antibody | Species | Dilution | Company (Catalog #) |
|---|---|---|---|
| anti-mouse ICAM1 | rat | 1:300 | abcam (ab119871) |
| anti-mouse collagen IV | rabbit | 1:300 | Bio-Rad (2150-1470) |
| anti-mouse F4/80 | rabbit | 1:300 | Cell Signaling (70076S) |
| anti-mouse Iba1 | rabbit | 1:300 | Fujifilm (019-19741) |
| anti-mouse Tmem119 | rabbit | 1:300 | Cell Signaling (90840S) |
| anti-rat IgG AF555 | goat | 1:300 | Invitrogen (A-21434) |
| anti-rabbit IgG AF647 | goat | 1:300 | Invitrogen (A-21244) |

#### Supplemental Figures

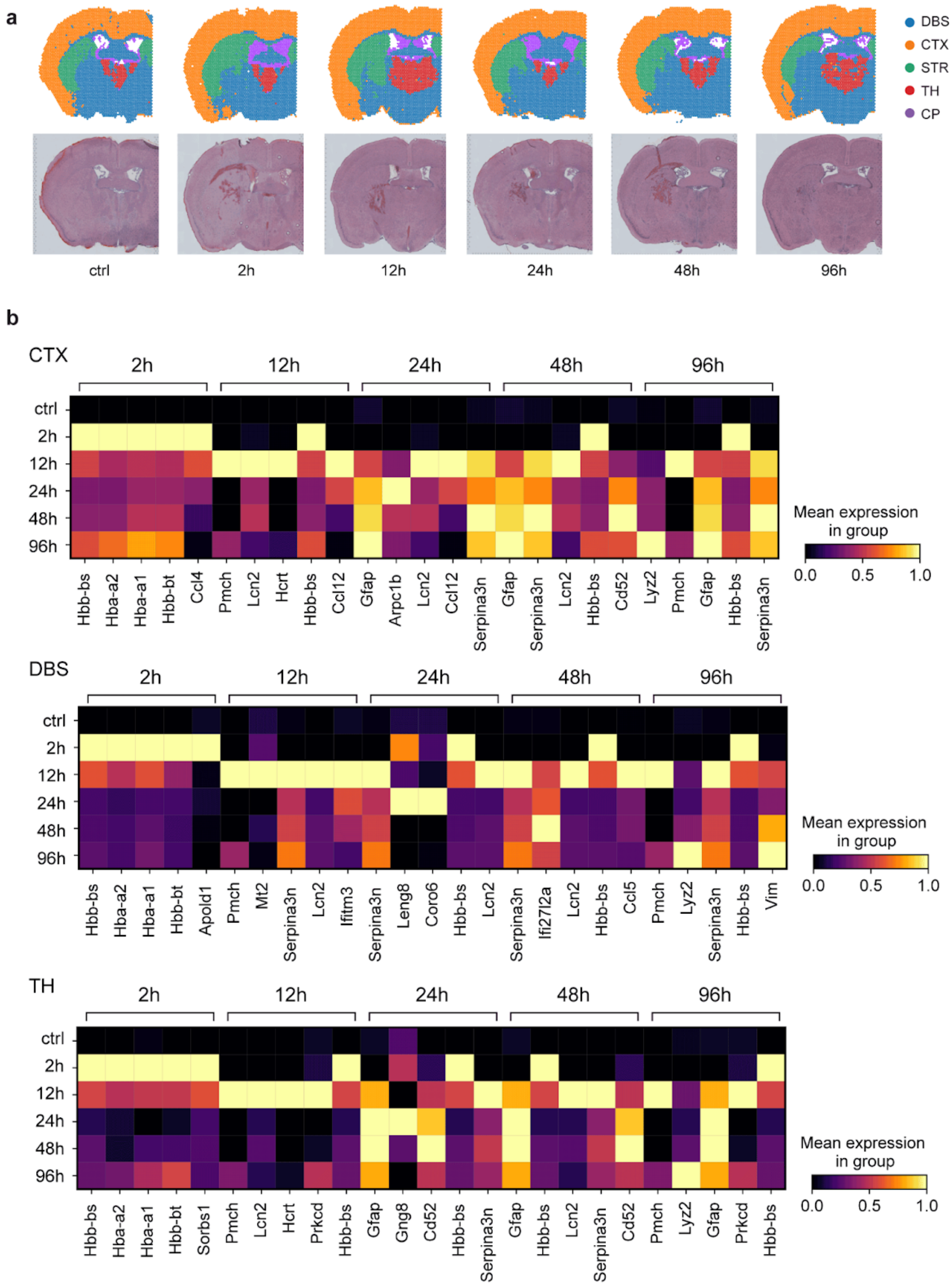


**Supplemental Fig 1. Spatial projection of unsupervised clusters.**

**A.** Spatial projection of features colored according to the allocated anatomical compartment and the underlying H&E-stained histological section for the control sample and samples at different time points after whole blood injection *(DBS, deep brain structures; CTX, cortex; STR, striatum; TH, thalamus; CP, choroid plexus).*
**B.** Top five differentially expressed genes (control as baseline) in the CTX (top), DBS (middle), and TH (bottom) stratified by time point after injection.

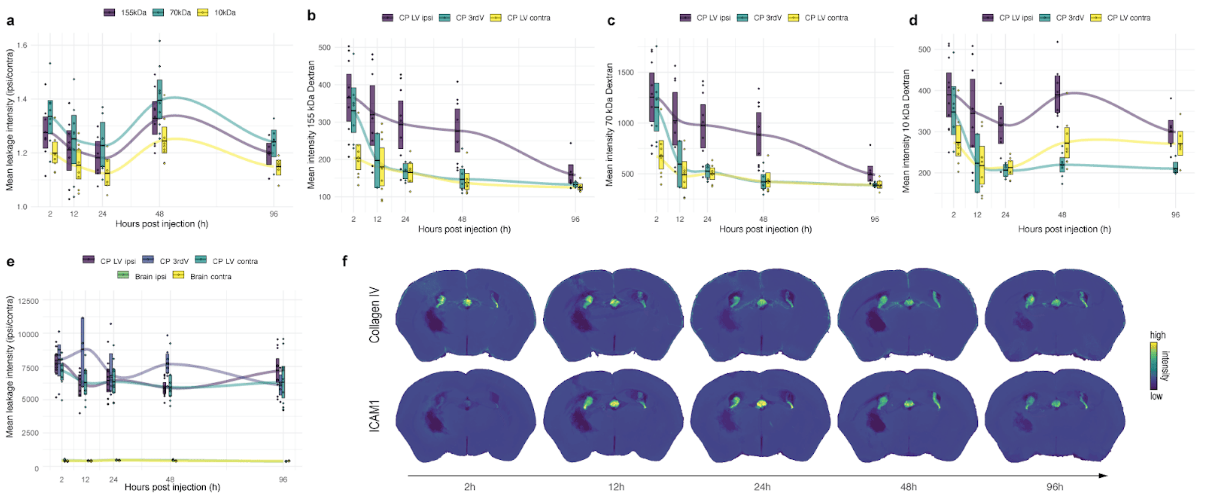


**Supplemental Fig 2. Temporal course of dextran perilesional extravasation, leukocytosis and ICAM1 signal intensity.**

**A.** Temporal dynamics of the normalized (ipsilateral/contralateral) mean fluorescence intensity for three dextran with different molecular weights (155 kDa, 70 kDa, 10 kDa) (n = 21).
**B.** Temporal dynamics of the mean fluorescence intensity of the TRITC labeled 155 kDa dextran in the different compartments of the choroid plexus (CP) (n = 21). *CP LV ipsi, choroid plexus ipsilateral to the injection site; CP 3rdV, choroid plexus of the third ventricle; CP LV contra, choroid plexus contralateral to the injection site.*
**C.** Temporal dynamics of the mean fluorescence intensity of the FITC labeled 70 kDa dextran in the different compartments of the CP (n = 21).
**D.** Temporal dynamics of the mean fluorescence intensity of the AlexaFluor647 labeled 10 kDa dextran in the different compartments of the CP (n = 21).
**E.** Temporal dynamics of the mean collagen IV signal intensity after striatal whole blood injection in different anatomical compartments (n = 21). *Brain ipsi, brain parenchyma ipsilateral to the injection site; Brain contra, brain parenchyma contralateral to the injection site.*
**F.** Average intensity maps for collagen IV (top row) and ICAM1 (bottom row) at different time points after striatal whole blood injection.
